# Degradation modeling of water environmental DNA: Experiments on multiple DNA sources in pond and seawater

**DOI:** 10.1101/2020.04.03.023283

**Authors:** Tatsuya Saito, Hideyuki Doi

**Affiliations:** Graduate School of Simulation Studies, University of Hyogo, 7-1-28 Minatojima-minamimachi, Chuo-ku, Kobe, 650-0047, Japan

**Keywords:** eDNA, decay rate, quantitative real-time PCR

## Abstract

Environmental DNA (eDNA) methods have been developed to detect organism distribution and abundance/biomass in various environments. eDNA degradation is critical for eDNA evaluation. However, the dynamics and mechanisms of eDNA degradation are largely unknown, especially when considering different eDNA sources, e.g., cells and fragmental DNA. We experimentally evaluated the degradation rates of eDNA derived from multiple sources, including fragmental DNA, free cells, and inhabiting species. We conducted the experiment with pond and seawater to evaluate the differences between freshwater and marine habitats. We quantified the eDNA copies of free cells, fragmental DNA, and inhabiting species (*Cyprinus carpio* in the pond and *Trachurus japonicus* in the sea). Our results show that eDNA derived from both cells and fragmental DNA decreased exponentially in both the sea and pond samples. The degradation of eDNA from inhabiting species showed similar behavior to the cell-derived eDNA. We evaluated three degradation models with different assumptions and degradation steps and found that a simple exponential model is effective in most cases. Our findings on cell- and fragmental DNA-derived eDNA provide fundamental information about the eDNA degradation process and can be applied to elucidate eDNA behavior in natural environments.

## INTRODUCTION

Environmental DNA (eDNA) methods are new methods developed for monitoring macroorganisms and managing aquatic ecosystems.^1–7^ eDNA is the DNA fragments released by organisms into an environment, such as water or soil. eDNA is thought to be derived from the feces,^8^ skin cells,^1^ mucus,^9^ and secretions^10^ of organisms. eDNA can be collected from environmental samples such as water^1^ and sediment.^11^ eDNA is mainly derived from fractions of cells or cellular organs but can also be derived from fragmental DNA in water.^12–13^

After the success of the first eDNA analysis, in which American bullfrogs were detected in French wetlands,^1^ the eDNA method was expanded for use with various aquatic taxa, including fish,^14^ reptiles,^15^ crustaceans,^16^ amphibians,^1^ aquatic insects,^17^ and mollusks^18–20^, and in various habitats, including ponds,^16,21^ rivers,^14,22,23^ lakes,^2,24^ and marine habitats.^25-27^ eDNA methods can be used not only to detect the target species, but also to estimate the biomass and abundance^2,23^ of the target species from the eDNA concentration. In just a decade, eDNA methods have rapidly developed and become a powerful tool for biomonitoring. Using eDNA methods without capturing individuals, biomonitoring surveys can be performed quickly and noninvasively.^16,21,28^

There are still many unclear points about the basic behavior of eDNA in water,^29^ especially the states and degradation of eDNA in the water^29-30^. Understanding eDNA states and degradation is essential for the effective sampling and storage of eDNA, and it would allow us to correctly interpret the results of species distribution and abundance/biomass estimations.

Many experiments have been conducted to reveal the detailed states and degradation rates of eDNA under various conditions.^31–35^ In most previous experiments, a target organism is placed in an aquarium and then removed. The eDNA is measured over an experimental period to evaluate eDNA release and degradation.^31–33^ In the most cases, the eDNA degradation curves declined exponentially and the degradation rate was very fast, often less than a week.^32–33^ In addition, water conditions, such as salinity,^36^ water temperature,^34–35^ and pH,^32,37^ influenced the eDNA degradation rate. Although eDNA can be derived from many sources, such as fragmental DNA and the DNA in free cells in the water,^12–13^ the current literature does not distinguish between the degradation rates of fragmental eDNA and free cell-derived eDNA. To understanding the details of eDNA degradation in nature, an experiment separating these eDNA sources is essential and has not yet been conducted.

The aim of this study is to separately observe and compare the degradation of fragmental eDNA and free cell-derived eDNA. In this experiment, we used water from the sea and a pond to confirm the effects of the water conditions, especially salinity, on degradation.^36^ From the results, we discuss the effective sampling and storage of eDNA and the correction of potential errors in the interpretation of eDNA results in field surveys.

## MATERIALS AND METHODS

### Experiment outline

We used sea, pond, and purified water (A300, AS ONE, Osaka, Japan) as DNA-free samples (Figure 1) and divided each into 12 bottles. Each bottle was added a solution of isolated cells (from *Oncorhynchus kisutch*) and fragmental DNA (from an internal positive control, IPC, Figure 1). The seawater and pond water contained the eDNA of Japanese jack mackerel (*Trachurus japonicus*) and common carp (*Cyprinus carpio*), respectively. We used *O. kisutch* tissue for the isolated cells because the species was not distributed in the sampling region. We conducted the experiment for 7 days. A Sterivex filter (Merck Millipore, Burlington, MA USA) was used to filter 500-mL samples of water and 1.5 mL of the filtrate from each bottle was collected (Figure 1). After extracting eDNA from the filtrate and the Sterivex filter, the copy number of each type of DNA contained in the Sterivex samples and filtrate were estimated by quantitative real-time PCR (qPCR, Figure 1–5).

**Figure 1.**
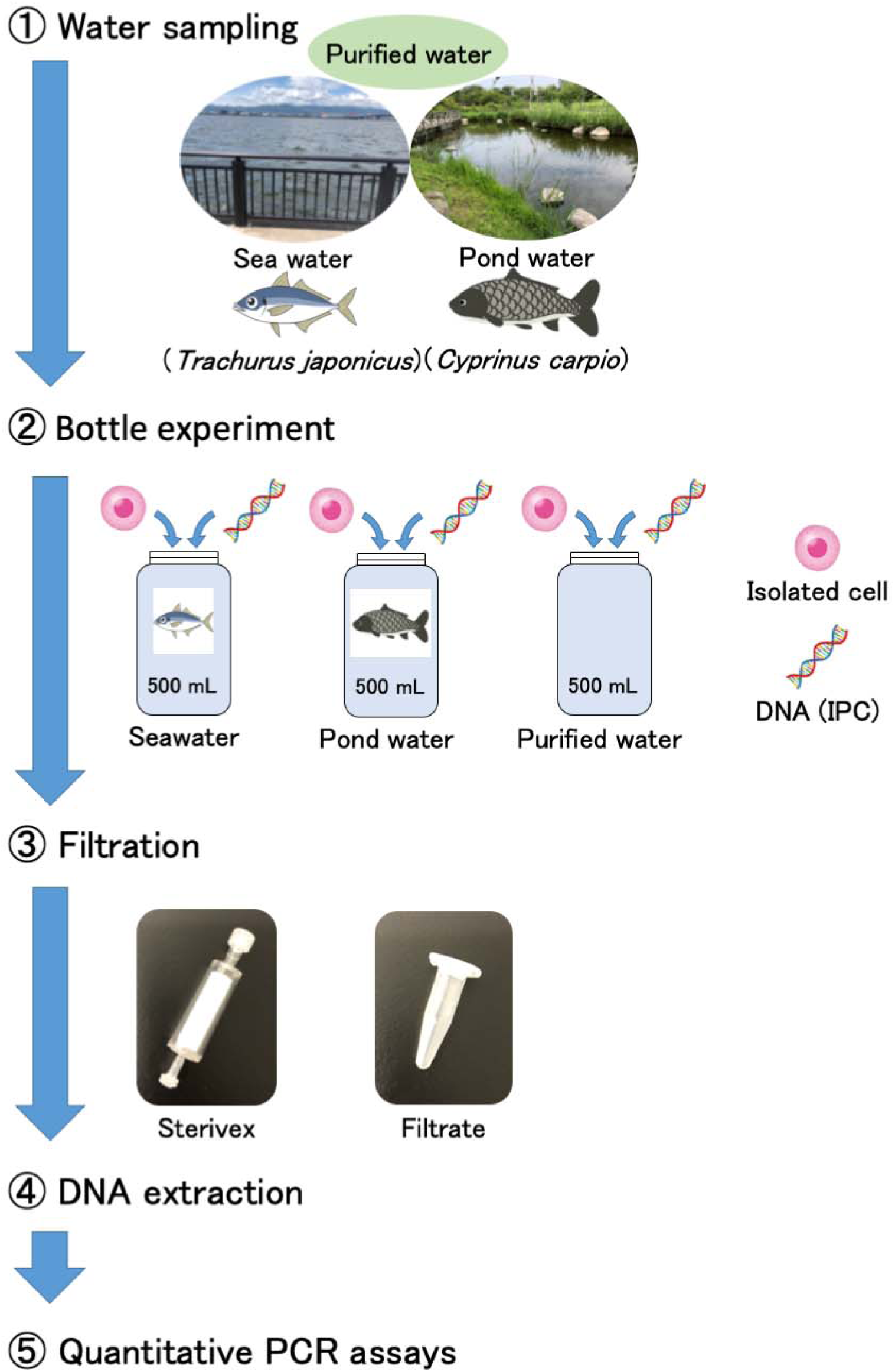
Experimental overview for the bottle experiments.

### Preparation of isolated cells

*O. kisutch* tissue was isolated using a Single Cell Isolation Kit (Cosmo Bio, Tokyo, Japan). We obtained the muscle of farmed *O. kisutch* in Chili. We placed 29-mg *O. kisutch* epidermal muscle samples onto the filter unit of the kit and added 100 µL cold buffer A, according to the manufacturer’s protocols. The tissue was then ground using a plastic rod 60 times. We added 400 μL buffer to the filter unit, then inverted it a few times and centrifuged it at 4000 rpm for 4 min. The filter unit was vortexed, then centrifuged at 2000 rpm for 5 min. We repeated the above procedure three times to prepare 1500 μL of isolated cells. The isolated cells were immediately used for the following experiments.

### Bottle experiment

We collected the sea and pond water from the Seto Inland Sea and an artificial pond in Kobe, Japan (34°38’28”N 135°13’37”E and 34°39’40”N 135°13’02”E, respectively) using bleached tanks. We bought the purified water for the experiment. The sea, pond, and purified water were divided into 12 bottles each. eDNA was removed from the bottles and equipment using 10% commercial bleach (ca. 0.6% hypochlorous acid) and washing with DNA-free distilled water. We set the bottles in a laboratory with around 25 °C room temperature at noon.

Each bottle received 100 μL of a solution of isolated cells and DNA [IPC, 207-bp (Nippon Gene, Tokyo, Japan); 1.5 × 10^5^ copies]. We collected and filtered 500 mL of water from each bottle using 0.45-µm Sterivex filters 0, 0.5, 1, 1.5, 3, 6, 12, 18, 24 (day 1), 48 (day 2), 72 (day 3), 120 (day 5), and 168 (day 7) h after the introduction of the cells and DNA, and collected 1.5 mL samples of the filtered water. After filtration, approximately 2 mL of RNAlater (Thermo Fisher Scientific, Waltham, MA USA) was injected into the Sterivex. As a filtration blank, the 500-mL DNA-free water was filtered in the same manner after filtration of the samples to monitor cross-contamination. The Sterivex filters and filtrate were immediately stored at –20 °C until further analysis.

### DNA extraction

The Sterivex filter was extracted using a DNeasy Blood and Tissue Kit (Qiagen, Hilden, Germany) following Miya et al.^38^ The RNAlater was removed using a 50-mL syringe, and 440 μL of the mixture [220 μL of phosphate-buffered saline (PBS), 200 μL of buffer AL, and 20 μL of proteinase K (Qiagen, Hilden, Germany)] was added to the Sterivex filter. We incubated the filters on a rotary shaker (at 20 rpm) at 56 □ for 20 min. We transferred the incubated mixture into a new 1.5 mL tube by centrifugation at 5000 g for 5 min. We then purified it using the DNeasy Blood and Tissue Kit, and finally eluted the DNA in 100 μL of buffer AE from the kit.

The filtrate was extracted using a DNeasy Blood and Tissue Kit, according to Uchii et al.^39^ We added 440 μL of the mixture (220 μL of PBS, 200 μL of buffer AL, and 20 μL of proteinase K) to 500 μL of the filtrate. We incubated it at 56 °C for 30 min, then added 200 μL of ethanol to the mixture and vortexed it for 2–3 s. We transferred the mixture into a new 1.5 mL tube by centrifugation at 5000 g for 5 min. We then purified it using the DNeasy Blood and Tissue Kit, and finally eluted the DNA in 100 μL of buffer AE from the kit. The extracted DNA from both methods was stored at –20 °C until qPCR analysis.

### Primer-probe design for O. kisutch

To detect and quantify the DNA of *O. kisutch* using qPCR, the forward and reverse primers for a 120-bp fragment of the COI region of the mitochondrial DNA were prepared from Chalde et al.^40^ and we designed TaqMan probe using Primer3Plus (http://www.bioinformatics.nl/primer3plus). The sequences of the real-time PCR primers and TaqMan probes are shown in Table 1.

**Table 1.**
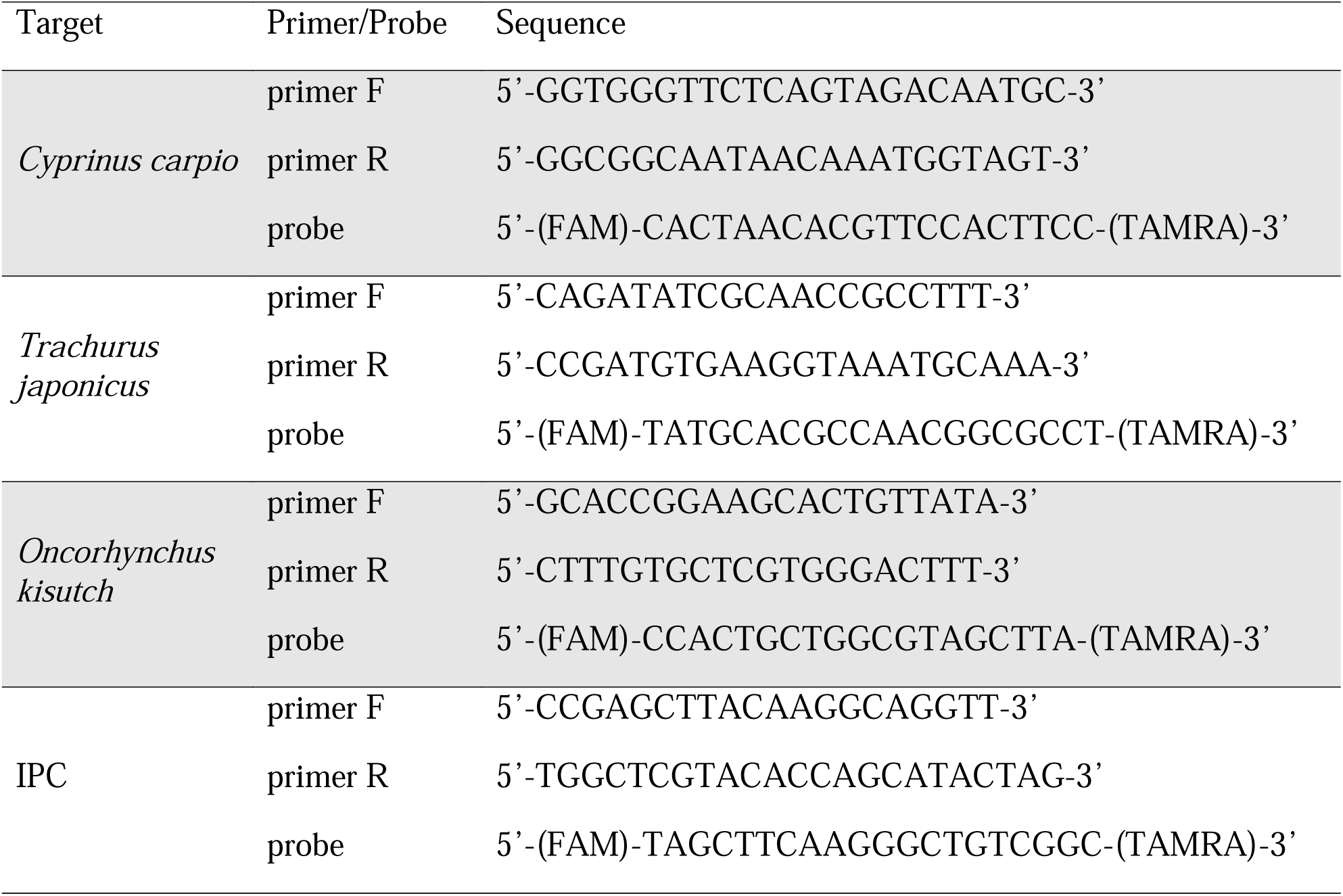
The primers and probes for the targeted DNA used in the experiment.

The specificity of the primers and probes was checked *in silico* using homologous sequences from other *Oncorhynchus* that inhabit Japan (National Center for Biotechnology Information, NCBI; http://www.ncbi.nlm.nih.gov/). No other *Oncorhynchus* genus were detected during the *in silico* screening for specificity, which was performed using Primer-BLAST (http://www.ncbi.nlm.nih.gov/tools/primer-blast/).

### Quantitative PCR assays

Quantitative PCR for *C. carpio*,^2^ *T. japonicus*,^41^ *O. kisutch*, and the IPC was performed. The DNA concentration was quantified by qPCR using a PikoReal™ qPCR system (Thermo Fisher Scientific, Waltham, MA, USA). Each TaqMan reaction contained 900 nM of forward and reverse primers and 125 nM of TaqMan probe in 1× TaqPath™ qPCR master mix (Thermo Fisher Scientific), and 2 µL of DNA sample was added to reach a final volume of 10 μL. Dilution series containing 1.5 × 10^1^ to 1.5 × 10^4^ copies per PCR plate were prepared and used as quantification standards. DNA of partial targets were cloned into plasmids and amplified in four replicates to obtain a standard curve. Quantitative PCR was performed with the following conditions: 2 min at 50 °C, 10 min at 95 °C, and 55 cycles of 15 s at 95 °C and 1 min at 60 °C). Four replicates were performed for each sample, and four replicate negative non-template controls containing DNA-free water instead of template DNA were included in all PCR plates. We performed the real-time PCR procedures according to the MIQE checklist. The PCR and qPCR were set up in two separate rooms to avoid the PCR-amplicon contamination.

The qPCR results were analyzed using PikoReal software ver. 2.2.248.601 (Thermo Fisher Scientific, Waltham, MA USA). A standard curve for the target was constructed using a dilution series of 15000, 1500, 150, and 15 copies per PCR reaction. For the standard curve, we used target DNA cloned into a plasmid. The R^2^ values of the standard curves ranged from 0.989– 0.998 and the PCR efficiency from 91.07–101.68%. The concentration of DNA in the water collected (DNA copies mL^−1^) was calculated from the volume of filtered water. DNA copy numbers were evaluated including negative amplifications as zero values. We evaluated the limit of detection (LOD) of the qPCR with four replicates for all the primers/probes used to detect *C. carpio, T. japonicus, O. kisutch*, and the IPC.

### Statistical analysis

All statistical analysis and data plotting were performed using R version 3.6.0 (R Core Team, 2019). Degradation models were established for determining a first-order rate constant from biotransformation/degradation studies, including standard biotic studies conducted in soil, water, or mixed media.^42–43^ The degradation rates in this study were estimated from the DNA degradation curves obtained from three models: the Single First-Order rate model (SFO), First-Order Multi-Compartment model (FOMC), and Double First-Order in Parallel model (DFOP).^42–43^ The SFO establishes a simple procedure for determining a first-order rate constant from the degradation of DNA. The FOMC establishes a procedure for determining how fast the degradation rate declines with decreasing concentration owing to the degradation of DNA, as well as determining a first-order rate constant from the degradation of DNA. The DFOP established a procedure for determining two first-order rate constants from the degradation of DNA.

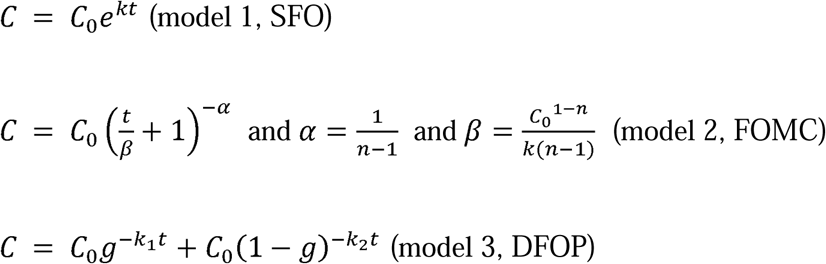

where *C* is the eDNA concentration at time t, *C*_0_ is the eDNA concentration at time 0, i.e., the initial eDNA concentration, and k is the degradation rate constant per hour. n determines how fast the degradation rate declines with decreasing concentration and is an indicator of how far the data deviate from a first-order model (where n = 1). lil, β, and, g are constants, which are estimated by analyzing the nonlinear least □ Jsquares regression. We performed all modeling using the “mkin” package version 0.9.49.8 in R. We evaluated the fitting of the models by chi-squared error level. Due to the differences in the model structures, we did not evaluate the differences using information criteria, such as AIC or BIC, which are generally used to evaluate the number of model parameters. Significant differences in model coefficients were evaluated by overlapping the 95% confidential intervals (CIs) of the coefficients (i.e., 11 = 0.05).

## RESULTS

### Primers and probe testing, and LOD

We confirmed that *O. kisutch* isolated cells were amplified using the primers and designed probe. We found the LODs of the qPCR were one copy per PCR reaction for all the primers/probes used to detect *C. carpio, T. japonicus, O. kisutch*, and the IPC.

### Sterivex

We detected all the targeted DNA of *C. carpio, T. japonicus*, the *O. kisutch* cells, and the IPC by qPCR (Figure 2). While we could not detect the eDNA of *C. carpio* and *T. japonicus* on day 2, we detected the IPC DNA and *O. kisutch* cells in the purified water and seawater up to day 7, and the IPC DNA and *O. kisutch* cells in pond water up to day 5 and 3, respectively. The DNA concentrations of the cells and IPC DNA decreased exponentially after adding them (day 0). We observed these trends in both the sea and pond samples, while they were not observed in the samples with purified water.

**Figure 2.**
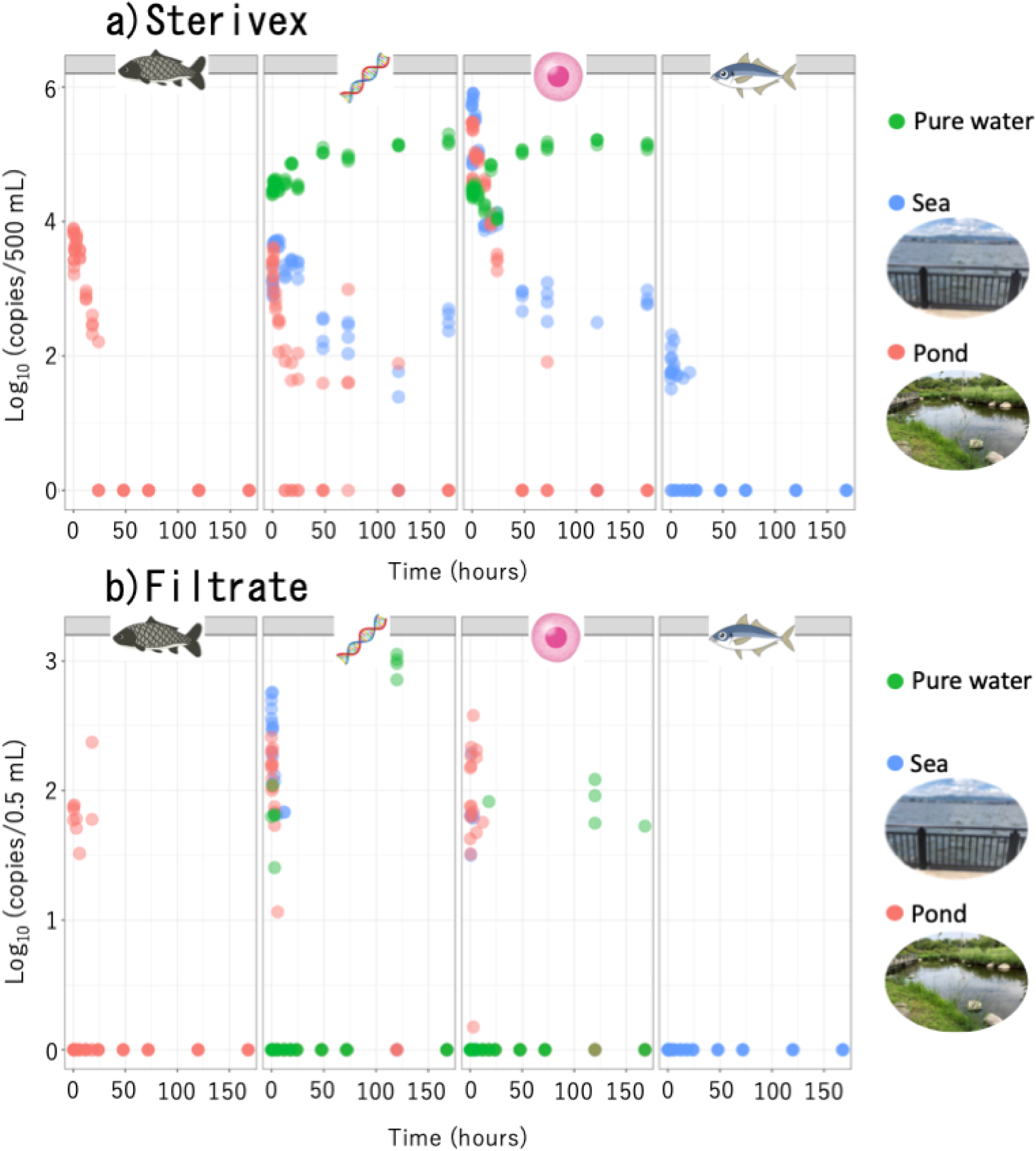
The relationship between the eDNA concentrations of the targets (*Cyprinus carpio*, the internal positive control, *Oncorhynchus kisutch* cells, and *Trachurus japonicus*), and the sampling timing of the experiment. The eDNA were extracted from a) the Sterivex and b) the filtrate. The dots indicate the eDNA concentrations of the targets at each time point under different water conditions: purified water, green; sea, blue; pond, red (N = 4 for each time point).

The SFO model was the most suitable model among the three degradation models after comparing their chi-squared error levels (Table 2). The SFO model showed the lowest chi-squared error levels between the three models (SFO, FOMC, and DFOP), except for a few samples. This result indicates that the efficiency of environmental DNA degradation did not decrease with time. However, the *O. kisutch* cells and *C. carpio* in the pond water had slightly lower chi-squared error levels for the DFOP than for the other two models. This result shows that the *O. kisutch* cells and *C. carpio* in the pond water had two different degradation rates: a faster rate was observed during the early part of the degradation curve and a slower rate was observed later on. The degradation rate constant of the IPC was significantly different between the sea and pond samples when comparing the 95% CIs (Figure 3). This indicates that pond–sea differences significantly affected eDNA degradation rate. The degradation constants of the *O. kisutch* cells, *C. carpio*, and *T. japonicus* were not significantly different between the sea and pond samples (Figure 3). The degradation constants of *C. carpio* and *T. japonicus* were not significantly different to that of the *O. kisutch* cells but were significantly different to that of the IPC (Figure 3).

**Table 2.**
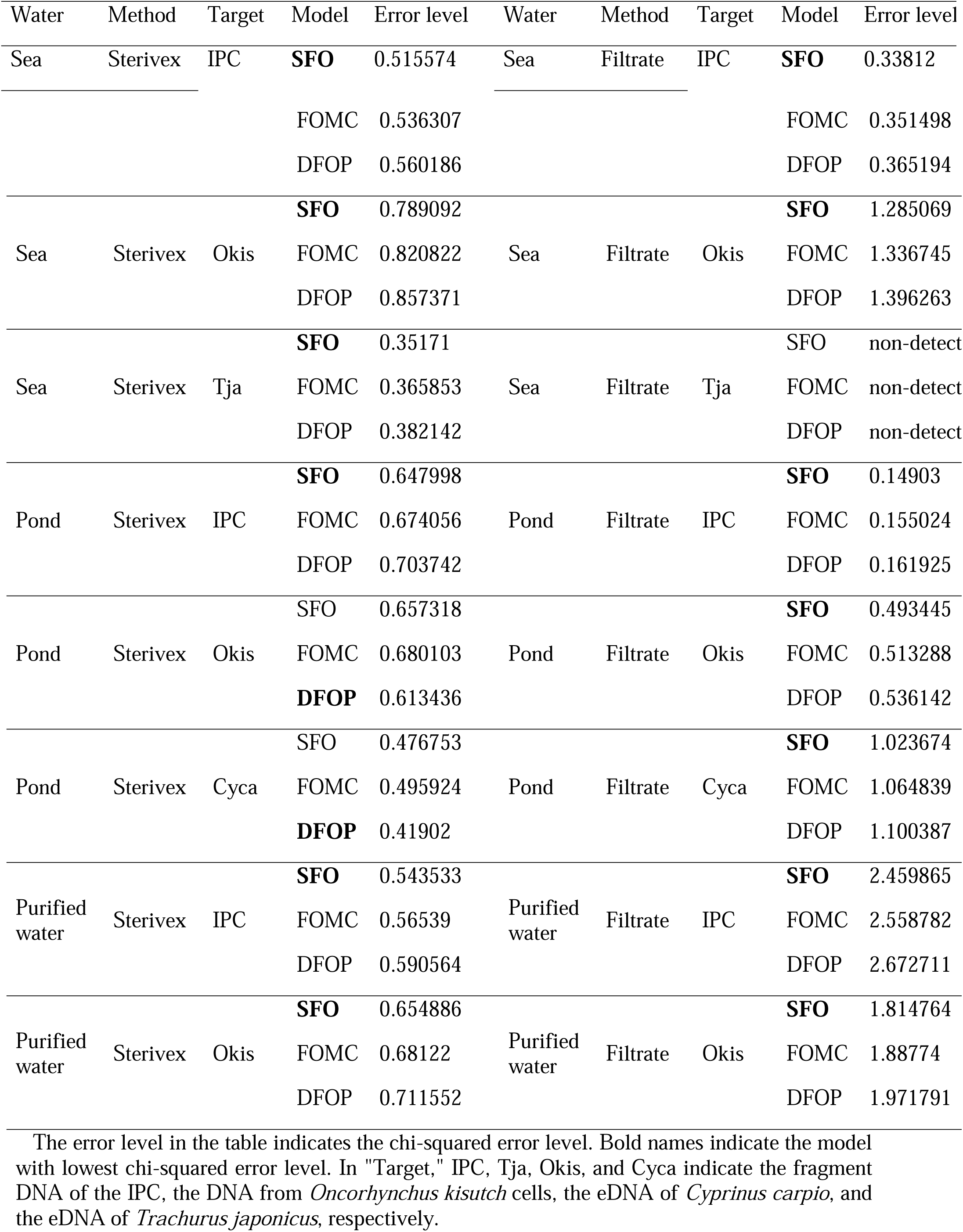
The fitting of the three models (SFO, FOMC, and DFOP) by chi-squared error level.

**Figure 3.**
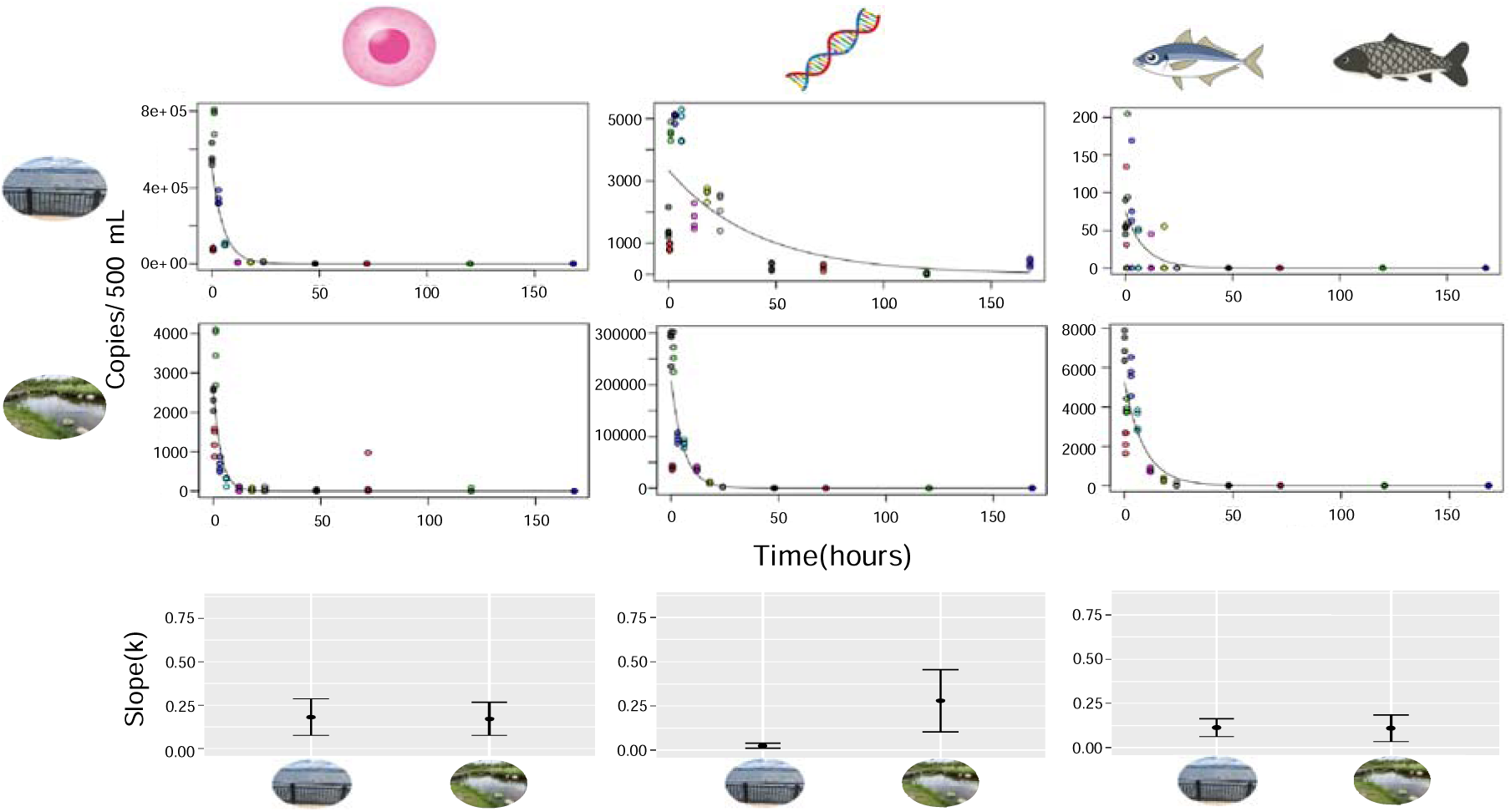
Degradation curves of the SFO and the rate constant for the bottle experiments using the Sterivex. The dots indicate the eDNA concentrations of the targets (*Cyprinus carpio*, the IPC, *Oncorhynchus kisutch* cells, and *Trachurus japonicus*) at each time point with different colors (N = 4 for each time point). The upper degradation curves show each target (*O. kisutch* cells, the IPC, and *T. japonicus*) in sea samples. The lower decay curves show each target (*O. kisutch* cells, the IPC, and *C. carpio*) in pond samples. The slope (k) of each target (*O. kisutch* cells, the IPC, *C. carpio*, and *T. japonicus*) is shown with 95% confidential intervals.

### Filtrate

We detected three of the targeted DNA, *C. carpio, O. kisutch* cells, and the IPC, by qPCR (Figure 2). However, we could not detect the IPC, *O. kisutch* cells, and the eDNA of *C. carpio* on day 2 in both the sea and pond. In the purified water, only the DNA of the IPC and *O. kisutch* cells were detected on day 5. We could not detect the eDNA of *T. japonicus* in any of the filtrate samples. The DNA concentrations of the cells and IPC decreased exponentially just after adding them (Figure 2). We observed these trends in both the sea and pond samples, while they were not observed in the samples with purified water.

The SFO model was also the most suitable for modeling the degradation of filtrate, except for a few samples (Table 2), indicating that the efficiency of environmental DNA degradation did not decrease with time. The degradation constant of the IPC was not significantly different between the sea and pond samples, while the degradation constant of *O. kisutch* cells was significantly different between the sea and pond samples (Figure 4).

**Figure 4.**
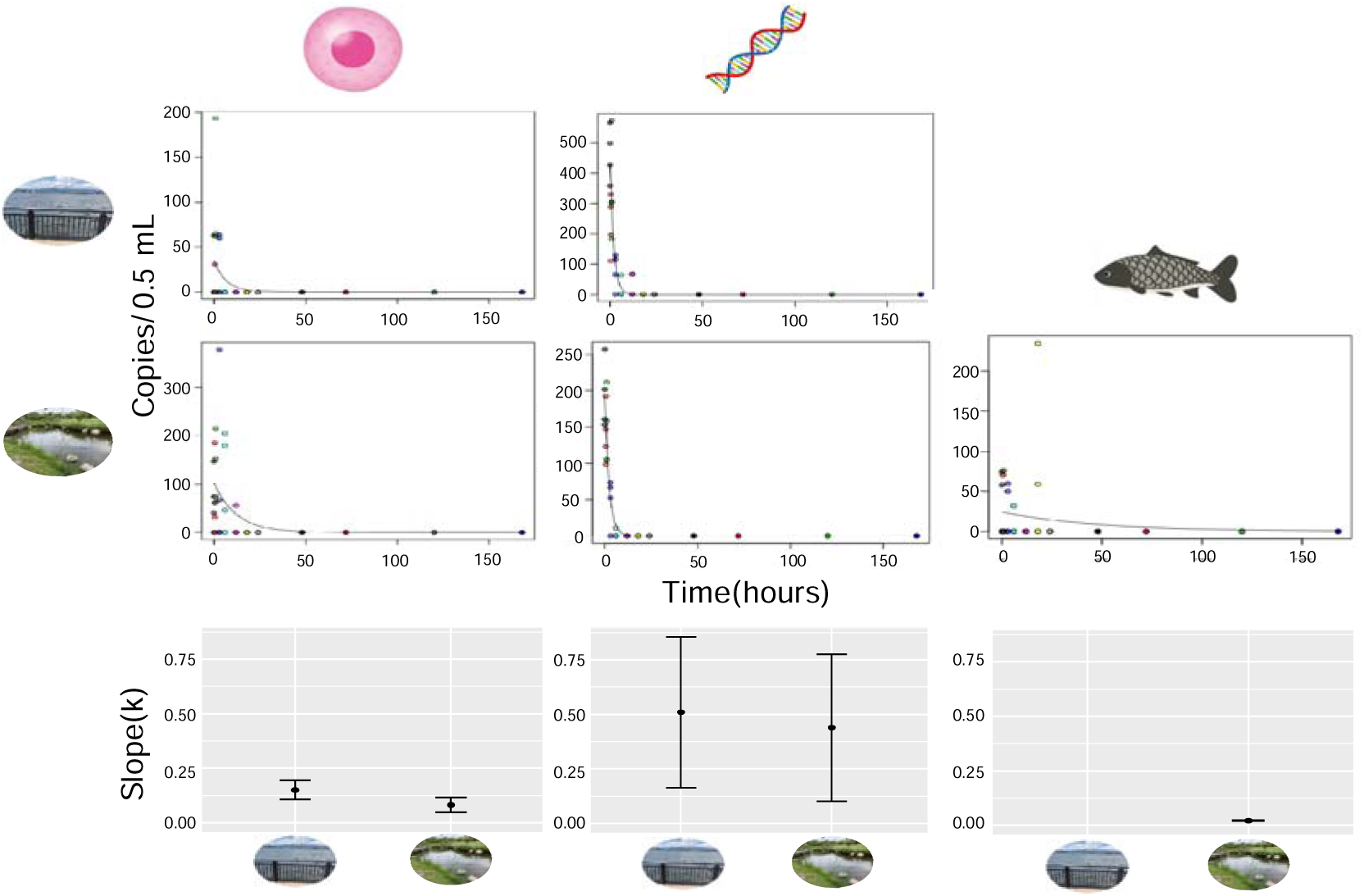
Degradation curves of SFO and the rate constant for the bottle experiments using the filtrate. The dots indicate the eDNA concentrations of the targets (*Cyprinus carpio*, the IPC, *Oncorhynchus kisutch* cells, and *Trachurus japonicus*) at each time point with different colors (N = 4 for each time point). The upper degradation curves show each target (*O. kisutch* cells and the IPC) in the sea samples. The lower decay curves show each target (*O. kisutch* cells, the IPC, and *carpio*) in the pond samples. The slope (k) of each target (*O. kisutch* cells, the IPC, and *C. carpio*) is shown with 95% confidential intervals.

## DISCUSSION

We found that the DNA concentrations of the *O. kisutch* cells and IPC declined exponentially in both the sea and pond water, although these DNA did not decline in the purified water. Our results of exponential degradation support those of previous studies.^2,33,34,35^ In purified water, the DNA concentrations did not decrease over time. eDNA degradation was mainly caused by microbes and extracellular enzymes,^29^ as well as UV radiation.^44^ Because the purified water did not contain microbes and extracellular enzymes and we performed the experiment in the laboratory, the DNA from the cells and the fragmental DNA in the purified water did not decrease over time. This supports the hypothesis that eDNA declines owing to microbes and extracellular enzymes rather than degrading by itself or through UV degradation.

We evaluated eDNA degradation rates using three models (the SFO, FOMC, and DFOP) to quantify general degradation processes. Previous studies estimated eDNA degradation rates by fitting simple exponential models (i.e., SFO),^31,34,45–48^ but did not compare the fit of multiple degradation models. Our results suggested that the SFO model can be used to evaluate eDNA degradation rate in most cases. The SFO model assumes that the degradation rate does not change over time. Thus, we can assume that the degradation processes in cell- and fragmental DNA-derived eDNA occur with similar timing, i. e., both cell decomposition and DNA degradation in the water occurred at almost the same rate in the experiments. However, the eDNA degradation of *O. kisutch* cells and *C. carpio* in the pond water fitted the DFOP better. The DFOP uses two different degradation rates in the model over time. Therefore, the eDNA degradation processes in the pond water might have two stages for the eDNA derived from cells and *C. carpio*. We speculate that there were two sequentially occurring degradation processes for the cell decomposition and free DNA degradation in the pond water. It is not known why two degradation processes were detected in the pond water but not in the seawater. Thus, further experiments are needed to reveal the details of this phenomenon.

The degradation rate of the eDNA derived from the inhabiting species, *C. carpio* and *T. japonicus*, in each site were not significantly different from that of the *O. kisutch* cells. This result might suggest that the degradation of the organisms’ eDNA in the water displays similar behavior to that of eDNA derived from free cells. Previous studies found that the most abundant eDNA size range was from 1 to 10 μm and concluded that eDNA is mainly derived from cells or cellular organs.^12–13^ Our findings may indirectly support the possibility that eDNA is mainly derived from cells. This result also suggests that the degradation rate of free cells can represent that of eDNA in nature, rather than the degradation rate of fragmental DNA. An experimental approach using free cells would be useful to experimentally reveal eDNA behavior in nature.

The fragmental DNA, the IPC, was degraded in the pond significantly faster than in the sea. Water salinity was found to be a significant factor in degradation rate and higher salinity sites had slower degradation rates in marine sites with salinity gradients.^36^ Our study supports these results of Collins et al.^36^ by experimentally comparing freshwater ponds and seawater. However, there were many differences in environmental factors between freshwater ponds and seawater, such as microbe abundance, species composition, and the other water quality properties. Further study is needed to confirm how the environmental factors in freshwater and marine habitats affect the degradation of eDNA derived from fragmental DNA.

Our experiments provide new findings on the eDNA degradation phenomenon; however there were some limitations owing to the experimental design. First, we performed the experiment using only one site each for the sea and pond samples. Therefore, it is unclear whether similar DNA degradation rates exist in other habitats. Experiments with various habitats need to be increased to understand eDNA degradation more generally. The evaluation of eDNA degradation while comparing different environmental conditions (e.g., salinity, water temperature, pH, chlorophyll, and microorganisms) may reveal what is affecting eDNA degradation in general. Second, we evaluated eDNA derived only from cells and fragmental DNA but other eDNA sources exist, such as cellular organs (e.g., mitochondria), cells with mucus, and tissue parts (e.g., skin, scales). Testing with these other sources would provide further information on eDNA degradation.

In conclusion, we found that the eDNA derived from cells and fragmental DNA declined exponentially just after being added to both sea and pond water samples. The eDNA from inhabiting species showed similar behavior to the eDNA derived from cells. A simple exponential model can, in most cases, be used to evaluate degradation. However, for cell-derived eDNA degradation freshwater ponds, we should consider the possibility of multiple degradation steps, such as cell decomposition and DNA degradation. A greater understanding and the accumulation of basic information about eDNA would improve eDNA methods and enable researchers to maximize the potential of future eDNA methods.

## Supporting information

https://www.icloud.com/numbers/0kU0lacWX6ZBCGPhGd7x1-3nQ#Table_S1

## ASSOCIATED CONTENT

### Supporting Information

The following files are available free of charge.

Table S1: All qPCR data (CSV file)

## AUTHOR INFORMATION

### Author Contributions

TS and HD designed the study, TS performed the experiments, TS contributed to the molecular experiments, TS and HD analyzed the data and interpreted the results, and TS and HD wrote the manuscript.

## ACKNOWLEDGMENTS

This study was supported by the Environment Research and Technology Development Fund (4-1602) of the Environmental Restoration and Conservation Agency, Japan and JST-CREST (JPMJCR13A2).

